# Supercoiling DNA optically

**DOI:** 10.1101/620005

**Authors:** Graeme A. King, Federica Burla, Erwin J. G. Peterman, Gijs J.L. Wuite

## Abstract

Cellular DNA is regularly subject to torsional stress during genomic processes, such as transcription and replication, resulting in a range of supercoiled DNA structures.^1,2,3^ For this reason, methods to prepare and study supercoiled DNA at the single-molecule level are widely used, including magnetic,^4,5,6^ angular-optical,^7,8,9^ micro-pipette,^10^ and magneto-optical tweezers.^11^ However, in order to address many open questions, there is a growing need for new techniques that can combine rapid torque control with both spatial manipulation and fluorescence microscopy. Here we present a single-molecule assay that can rapidly and controllably generate negatively supercoiled DNA using a standard dual-trap optical tweezers instrument. Supercoiled DNA formed in this way is amenable to fast buffer exchange, and can be interrogated with both force spectroscopy and fluorescence imaging of the whole DNA. We establish this method as a powerful platform to study both the biophysical properties and biological interactions of negatively supercoiled DNA.

## Introduction

Changes in both the twist and writhe of DNA play a significant role in many aspects of genome biology.^1, 2, 3^ The topology of DNA is defined by its linking number (*Lk*), which is a linear combination of the molecular twist and writhe. The linking number of DNA is typically tuned *in vivo* through torsional stress. The resulting change in topology is known as supercoiling (σ), and is quantified as the fractional change in *Lk, i.e*., (*Lk*-*Lk*_0_)/*Lk*_0_. Supercoiling can be either positive (σ > 0) or negative (σ < 0) depending on the direction of the applied torque. While both positive and negative supercoiling occur frequently *in vivo*, overall the genome of Prokaryotes and Eukaryotes is slightly negatively supercoiled.^2,12^ At very low DNA tensions (< 1 pN), torsional stress is predominantly absorbed as writhe, yielding buckled structures known as plectonemes.^13^ At slightly higher tensions (> 1 pN), negative supercoiling can induce a range of underwound DNA conformations.^4,6,9,14^ The ability to study the influence of supercoiling on DNA-protein interactions is therefore of great importance. However, despite recent technical advancements,^14,15,16,17^ it is still challenging to combine DNA torque control with several key functionalities, such as fluorescence imaging and rapid buffer exchange. To address this, we have developed a multi-functional nano-mechanical assay, termed Optical DNA Supercoiling (ODS).

## Results and Discussion

Our method exploits the intrinsic mechanical properties of DNA to induce a fixed reduction in *Lk* (Fig. 1a). Here, an end-closed DNA molecule is first rendered torsionally constrained (*i.e. Lk* is fixed) through the binding of at least two biotin moieties (on each end of the DNA) to streptavidin-coated, optically trapped beads.^18^ Application of high tension (~115 pN) induces overstretching, resulting in ~70% elongation in the DNA length without any global change in *Lk*.^18,19^ In the absence of torsional constraint, however, overstretching occurs at much lower forces (~65 pN),^20,21^ and is associated with cooperative unwinding of the double-helix; in this case, the average molecular twist is reduced from ~10.5 bp/turn to ~37.5 bp/turn.^19^ Consequently, when torsionally constrained DNA is overstretched, the molecule experiences torsional stress, relative to the unconstrained state. This torsional stress can be released through swivelling of the DNA around a single biotin-streptavidin tether, which can happen for example when one of the biotin-streptavidin connections is disrupted (Fig. 1b). If the disrupted tether subsequently reforms, torsional constraint will be reinstated, but now with the DNA in a lower *Lk* than that of the B-form double-helix (Fig. 1a). Crucially, the reduced *Lk* is preserved upon decreasing the tension (*i.e*. the DNA remains negatively supercoiled).

**Figure 1.**
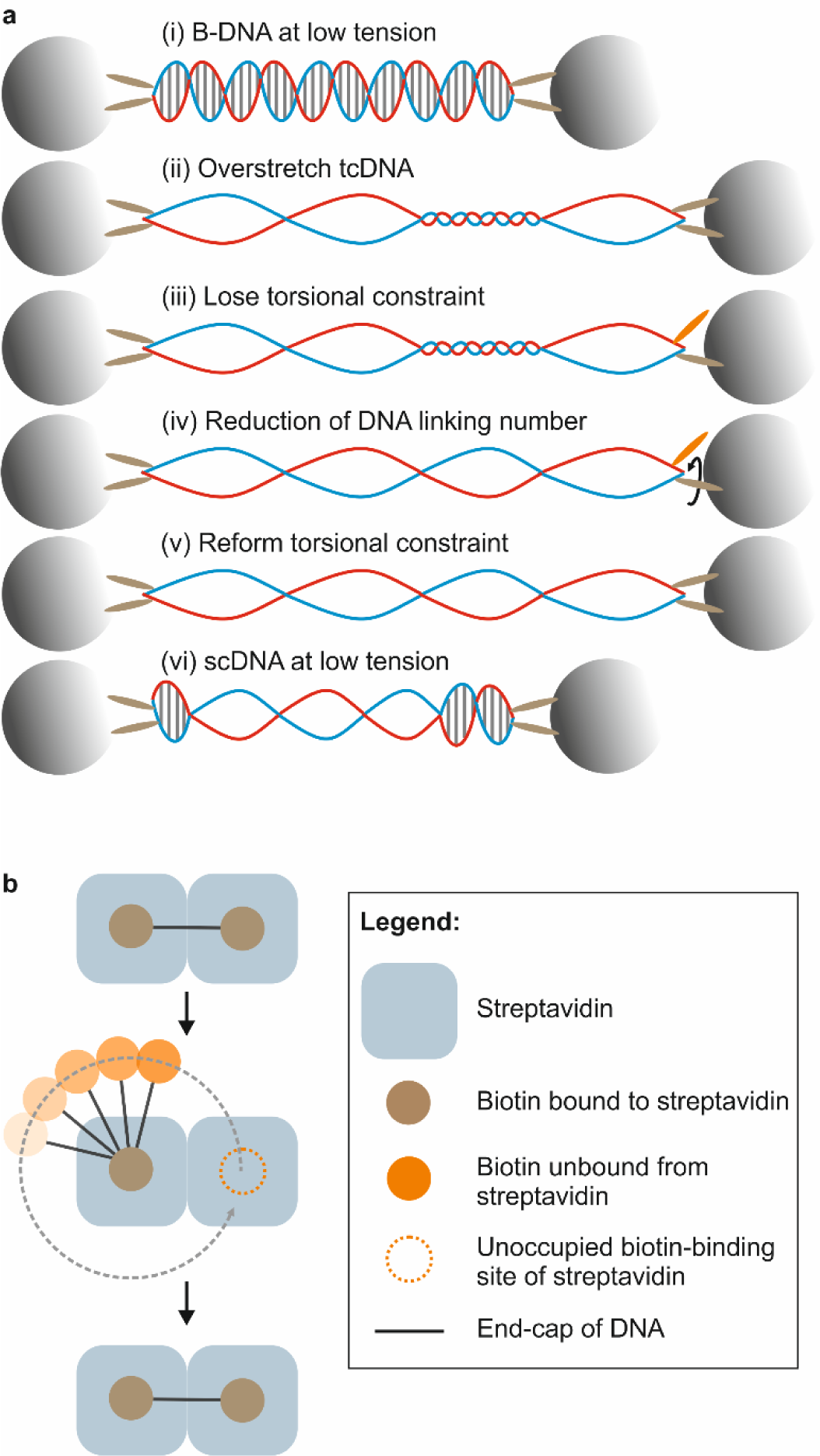
Generation of negatively supercoiled DNA using dual-trap optical tweezers. **a**. (i) End-closed DNA is torsionally constrained between two optically trapped beads via at least two biotin-streptavidin bonds (brown ellipses) on each end of the molecule. (ii) The torsionally constrained DNA molecule (tcDNA) is overstretched via displacement of one of the beads. (iii) At high force (at least ~80 pN), a sufficient number of biotin-streptavidin bonds break (orange ellipse) such that only a single tether is present on at least one end of the DNA molecule. This results in the loss of torsional constraint. (iv) The DNA molecule unwinds by swivelling around the single tether (arrow). (v). Once the linking number has decreased, the broken biotin-streptavidin bond(s) can reform. The molecule is once again torsionally constrained, but now in a lower linking number than that of B-DNA. (vi) The tension is released by reducing the DNA extension, stabilizing the negatively supercoiled (sc) state. **b**. Schematic illustration of the tethering geometry that results in the formation of constrained underwound DNA. One biotin moiety (attached to the end-cap of DNA) transiently unbinds from streptavidin at high force, during which time a second biotin remains bound to another streptavidin unit. The dashed grey arrow represents DNA unwinding during the time that the biotin-streptavidin bond is disrupted.

We demonstrate this process experimentally in Fig. 2. Here, the sequential force-distance curves obtained (following overstretching of end-closed torsionally constrained DNA) correspond to those expected for negatively supercoiled molecules (with values of σ ranging from ~-0.1 to ~-0.7) (Fig. 2a).^19^ The magnitude of σ can be calibrated with reference to the known force-extension curves of negatively supercoiled DNA (Fig. 2b and Methods).^19^ The maximum value of σ that can be generated using ODS is ~-0.7: the same as that associated with fully overstretched unconstrained DNA.^19^ This value of σ will occur if the torsional stress is fully released by transient disruption of a biotin-streptavidin tether at high force. In most cases, however, the tether reforms before the limiting value of σ is reached. Importantly, the probability and timescale for tether disruption can be tuned by holding the torsionally constrained DNA molecule at high tension for increasing periods of time. For instance, after 10 seconds at ~150 pN, the average value of σ generated is typically < 0.1 (Fig. 2c). However, as demonstrated in Fig. 2a, by repeating such stretch-release cycles with the same DNA molecule, the value of σ can be tuned over a much wider range (up to σ ~-0.7). The above behaviour accounts for ~25 % of cases, where supercoiling is only generated at forces > 115 pN (*i.e*. beyond the end of the overstretching transition). In another 25-35% of molecules, one of the biotin-streptavidin tethers is even less stable, and will already transiently disrupt during overstretching – this allows the magnitude of σ to be even more precisely controlled, simply by tuning the extent to which the molecule is initially overstretched (Fig. 2d,e). A detailed summary of the various force-distance behaviours of end-capped torsionally constrained DNA is provided in Figs. S1 and S2 and SI Note 1.

**Figure 2.**
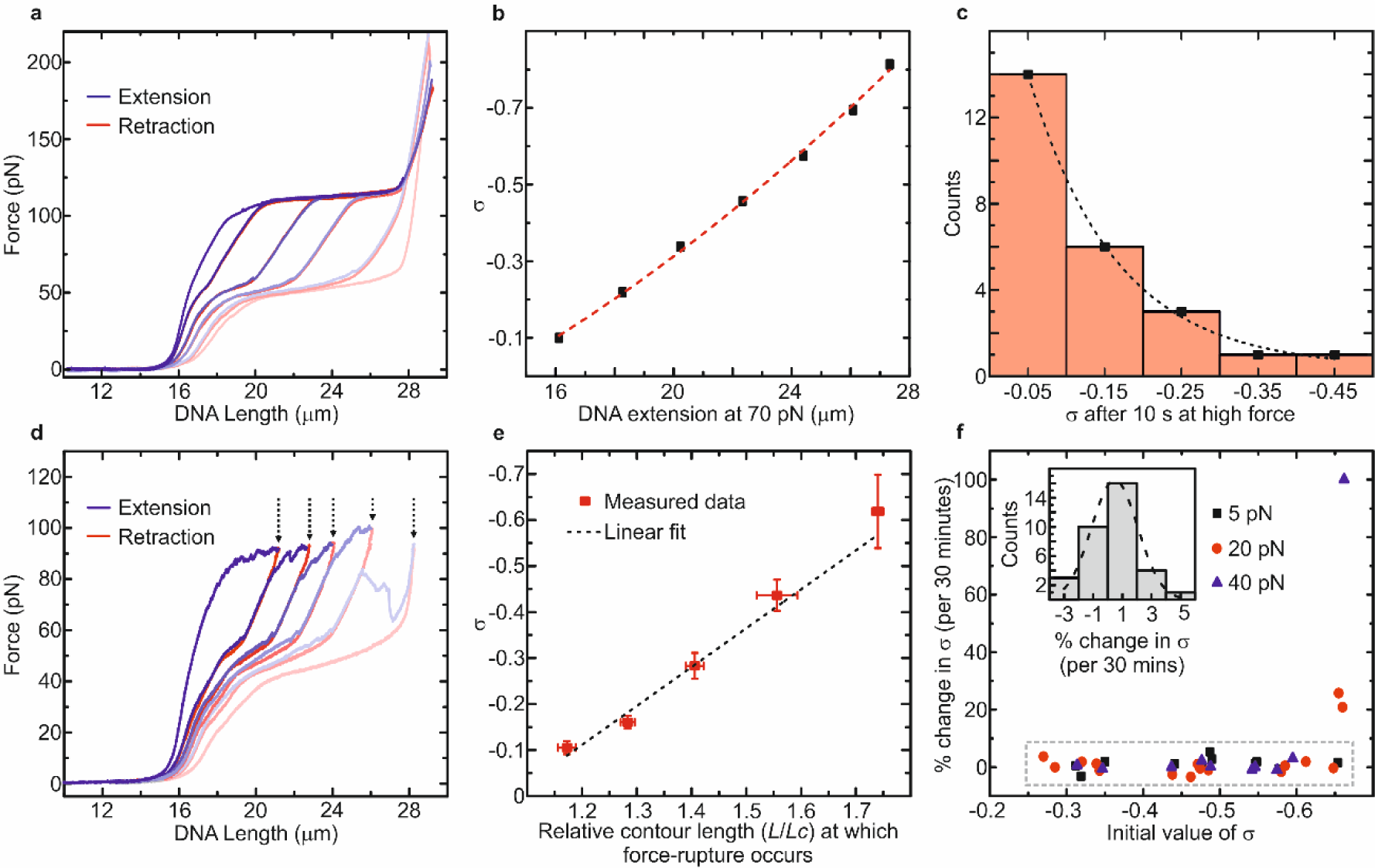
Controlling the magnitude and stability of the supercoiled state using tension. **a**. In a subset of molecules (~25%), the extent of supercoiling (σ) can be controlled by repeatedly extending the molecule (blue) and waiting at high forces (> 115 pN) for a given period of time. The resulting force-retraction traces (red) are consistent with those expected for negatively supercoiled DNA.^19^ Force-distance curves associated with repeated stretch/relax cycles are shown by the fading blue/red colour scheme. **b**. A reference plot showing the change in σ as a function of DNA extension at 70 pN, as determined from previously published force-extension curves of supercoiled DNA.^19^ These data are interpolated using a second order polynomial (dashed red line). **c**. Histogram showing that, for the subset of molecules that behave similar to those in panel a, the average value of σ generated after 10 seconds at high (~150 pN) force is ~-0.1 (*N* = 25). Note, though, that much larger magnitudes of σ (up to ~-0.7) can be generated by repeating the above procedure several times (as shown by the different force-distance curves in panel a). **d**. In another subset of molecules (~25-35%), the supercoiled state is generated following force ruptures that occur between 80 pN and 120 pN (blue). After each force rupture, an increasing magnitude of supercoiling is formed (as evidenced by the force-retraction curves, red). **e**. For the behaviour described in panel d, the average amount of supercoiling formed is roughly proportional to the extension at which the force rupture occurred (*N* = 15). **f**. The fractional change in σ over 30 minutes for supercoiled DNA held at a constant force (5 pN, 20 pN or 40 pN). Inset: histogram showing the change in σ for the data contained within the grey dashed box. All data were recorded in a buffer of 20 mM Tris-HCl, pH 7.6, with 25 mM NaCl. Errors are s.e.m.

As well as being able to control the magnitude of σ, we note that, once generated, the supercoiled state is stable (*i.e. Lk* is fixed) over long periods of time at low forces. This is attributed to two key features. First, the biotin-streptavidin bond is orders of magnitude stronger at lower forces (*e.g*. < 10 pN) than at higher forces (> 100 pN) and is thus significantly less likely to break at low force.^22^ Second, provided large microspheres are used, the torsional stress is not significantly dissipated through rotation of the beads (Fig. S3, SI Note 2). Experimentally, we verify that in ~90% of cases, the linking number of the supercoiled molecule changes < 2% over a period of 30 minutes at forces in the range of 5-40 pN when using 4.5 μm diameter beads (Fig. 2f, Fig. S4, SI Note 3).

Since ODS is based on a standard dual-trap optical tweezers assay, it is inherently compatible with a wide range of functionalities, not least imaging of the whole DNA molecule with fluorescence microscopy.^23,24^ Additionally, the supercoiled substrate can be freely moved between different micro-fluidic channels, allowing for fast buffer exchange. This latter feature is greatly advantageous because it facilitates sequential binding of different proteins to the substrate, and also enables fluorescence images to be recorded under background-free conditions.^23^ Furthermore, in our technique, the supercoiled state is generated rapidly (on the order of seconds), independent of the length of the DNA molecule. This is important, because it facilitates the use of much longer substrates (such as λ-DNA), which is beneficial for many fluorescence imaging studies.^23^

In order to demonstrate the power of these combined functionalities, we first apply ODS to study the structural and mechanical properties of negatively supercoiled DNA as a function of ionic strength, tension, and protein binding. It has been previously reported that upon changing σ from −0.1 to −1.5, torsionally constrained DNA held at ~5 pN undergoes a cooperative transition from B-DNA to an increasingly underwound state.^6, 9, 14^

The underwound regions have been reported to exist as several different structures, including left-handed conformations (L-DNA and Z-DNA) as well as less defined base-pair melted conformations (bubble-melted DNA).^14^ However, the exact structures of underwound DNA and their dependence on local (*e.g*. cellular) environment is unclear. To gain a better understanding of this, we investigated the mechanical properties of negatively supercoiled DNA (for σ ~−0.45 and −0.65) by recording force-extension curves as a function of ionic strength (Fig. 3a,b). These experiments reveal two notable features. The first is that, for NaCl concentrations below ~50 mM, the overstretching transition appears less cooperative (*i.e*. the change in force as a function of extension is more gradual and less smooth). Second, at very low ionic strength (< 25 mM NaCl) hysteresis is observed when comparing force-extension and force-retraction curves (Fig. 3b). A similar hysteresis is also induced at elevated temperature (Fig. S5). These observations indicate that underwound DNA exhibits a heterogeneous structure at these conditions, and is at least partially base-pair melted at low ionic strength. By fitting the extensible Worm-Like Chain model (up to 30 pN), quantitative information can be extracted from these curves (Figs. 3c and S6). These fits reveal a significant decrease in the apparent DNA persistence length, from ~50 nm to ~30 nm as σ is reduced from −0.1 to −0.7. Concomitant with this, the apparent DNA contour length increases by ~8 % over the same range of σ. This is consistent with the presence of both L-DNA and bubble-melted DNA, which have each been reported to exhibit a lower persistence length and a greater contour length than B-DNA.^9^

**Figure 3.**
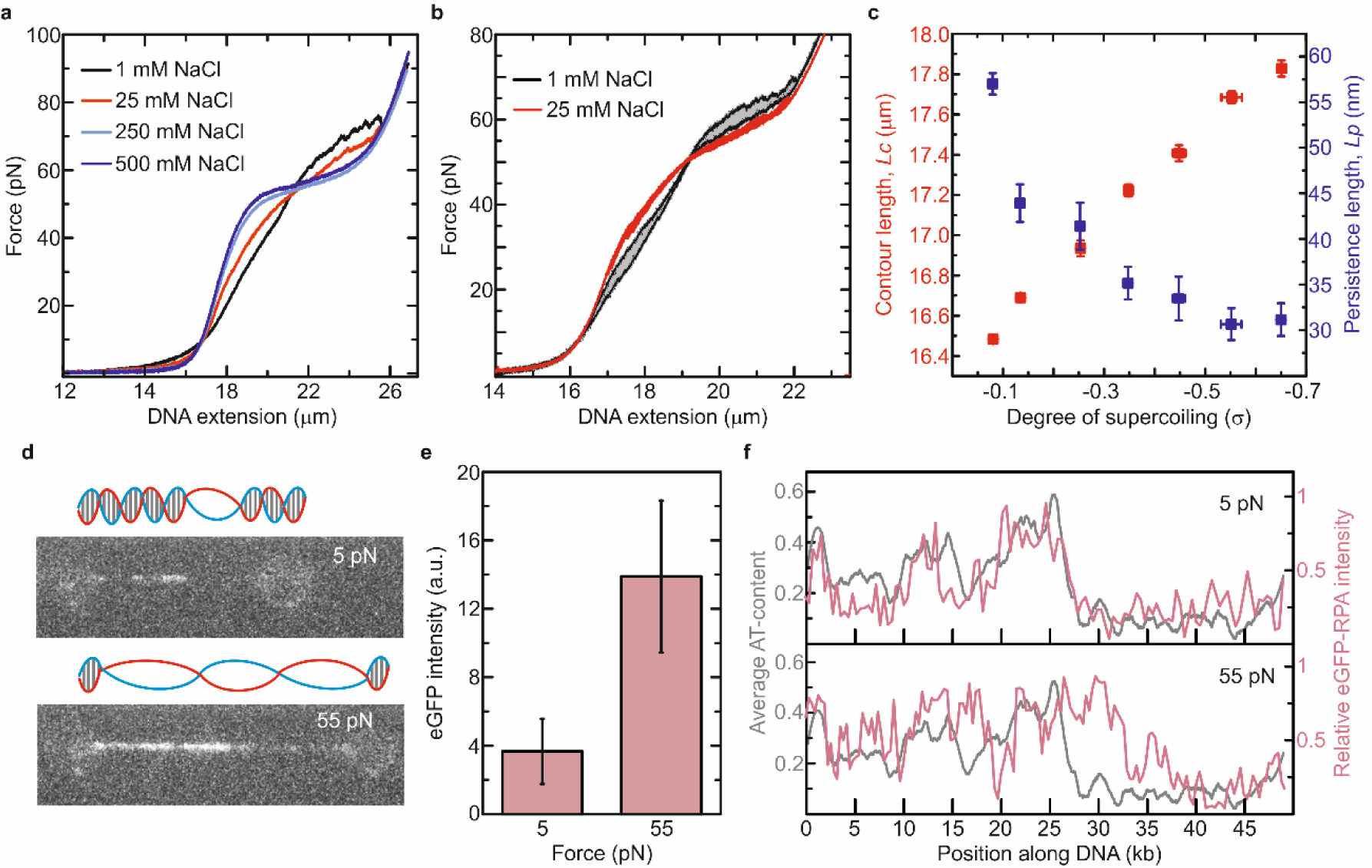
Application of ODS to study the structural and mechanical properties of underwound DNA. **a**. Force-extension curves of negatively supercoiled DNA (σ ~ −0.65) recorded in a range of NaCl concentrations (1 mM – 500 mM). **b**. Comparison of force-extension and force-retraction curves of negatively supercoiled DNA (σ ~ −0.45) in 1 mM NaCl and 25 mM NaCl, respectively. Filled grey areas highlight the observed hysteresis. **c**. Results of fits of the extensible Worm-Like Chain (eWLC) model to experimentally derived force-distance curves of end-closed DNA at 25 mM NaCl (up to 30 pN) as a function of supercoiling (s). Note that the DNA is expected to consist of regions of both B-DNA and underwound structures (such as L-DNA). Thus, the contour length (*Lc*) and persistence length (*Lp*) extracted from eWLC fits are average values for the entire DNA molecule, rather than for one specific structure. **d**. Sample fluorescence images for eGFP-RPA bound to negatively supercoiled DNA (σ ~ − 0.65) at 5 pN and 55 pN, respectively. Images were recorded in the presence of the protein (0.8 nM) after incubation for 2 minutes. **e**. Comparison of the average fluorescence intensity of eGFP-RPA bound to negatively supercoiled DNA (σ ~ −0.65) at 5 pN (*N* = 5) and 55 pN (*N* = 6). **f**. Comparison of the eGFP-RPA fluorescence intensity with the average AT-content along the length of the DNA molecule (at both 5 pN and 55 pN). All data were obtained in a buffer of 20 mM Tris-HCl, pH 7.6. Unless specified otherwise, the buffer contained 25 mM NaCl. Errors are s.e.m.

To gain more insight into the structure of underwound DNA, we imaged the binding of fluorescently-labelled Replication Protein A (eGFP-RPA) to negatively supercoiled DNA. This protein has previously been used to identify bubble-melted domains of DNA under mechanical strain.^14, 18, 20^ We first confirm that at low concentrations (0.8 nM), RPA does not significantly perturb the structure of underwound DNA (Fig. S7). Next, we compare the binding of eGFP-RPA to negatively supercoiled DNA at 5 pN and 55 pN, respectively (for σ ~-0.65). Note that the binding affinity of RPA for base-pair melted DNA is tension independent.^20^ These experiments were performed by incubating the DNA molecule in RPA (0.8 nM) for 2 minutes and then recording a snapshot image (*e.g*. Fig. 3d). The images reveal three important observations. The first is that eGFP-RPA binding is observed at low forces (~5 pN). This suggests that, at low tensions, localized regions of the DNA are base-pair melted and sufficiently accessible for RPA (which has a footprint of ~30 nucleotides^20^). This observation is consistent with our force-extension analysis (Fig. 3a,b and Fig. S5). The second key finding is that a significant (~3-4-fold) increase in RPA binding to negatively supercoiled DNA is observed at 55 pN, compared with 5 pN (Fig. 3d,e and Fig. S8), with force assisting the further melting of the DNA. Third, eGFP-RPA binding (at both low and high forces) is observed primarily in the most AT-rich domains of the DNA molecule (Fig. 3f). Such sequences have been shown previously to promote base-pair melting under mechanical stress.^20^ Together, the above findings support a growing body of evidence that negatively supercoiled DNA exhibits a heterogeneous structure at low forces (*e.g*. 5 pN) and low salt concentrations (*e.g*. 25 mM NaCl), with a co-existence of at least L-DNA, B-DNA and bubble-melted structures.^5, 6, 9, 14^

We next demonstrate the power of ODS to probe protein dynamics on supercoiled DNA. To this end, we investigate how negative supercoiling can influence the diffusion of the mitochondrial transcription factor TFAM (Mitochondrial Transcription Factor A). This protein has vital roles in transcription initiation, replication and transmission of mitochondrial DNA.^25,26^ TFAM has been shown previously to diffuse rapidly on B-form DNA via a 1D sliding motion, with a diffusion coefficient of ~10 × 10^4^ nm^2^ s^−1.26^ It has been proposed that this sliding provides a mechanism for the protein to efficiently search for its promoter sites on the mitochondrial genome.^26^ Notably, mitochondrial DNA is circular (and therefore torsionally constrained) and is believed to exhibit a range of topologies *in vivo*, including negatively supercoiled states.^27^ However, the influence of supercoiling on TFAM diffusion and on protein diffusion in general is unknown. To address this, we used ODS to probe three distinct DNA substrates, as highlighted in Fig. 4a: (i) B-form DNA at 5 pN, (ii) underwound DNA at 5 pN (σ ~-0.65) and (iii) overstretched unconstrained DNA (containing stretched bubble-melted structures). We then imaged monomers of alexa-555-TFAM (TFAM fluorescently-labelled with Alexa Fluor 555) on each DNA substrate, respectively (*e.g*. Fig. 4b). Using single-particle tracking, we plotted the average mean squared displacement (MSD) for TFAM monomers on each substrate over time (Fig. 4c). In all cases, the MSD increased linearly, consistent with free 1D diffusion. However, the diffusion coefficient for TFAM (determined from the gradient of the slopes in Fig. 4c) varies markedly for the different substrates. As summarized in Fig. 4d, the diffusion of TFAM on negatively supercoiled DNA (σ ~-0.65) at 5 pN is ~50% lower than on pure B-form DNA at the same force. Moreover, a further reduction in diffusion coefficient occurs on overstretched, unconstrained DNA. The above results indicate that TFAM mobility on DNA is highly sensitive to the presence of underwound structures.

**Figure 4.**
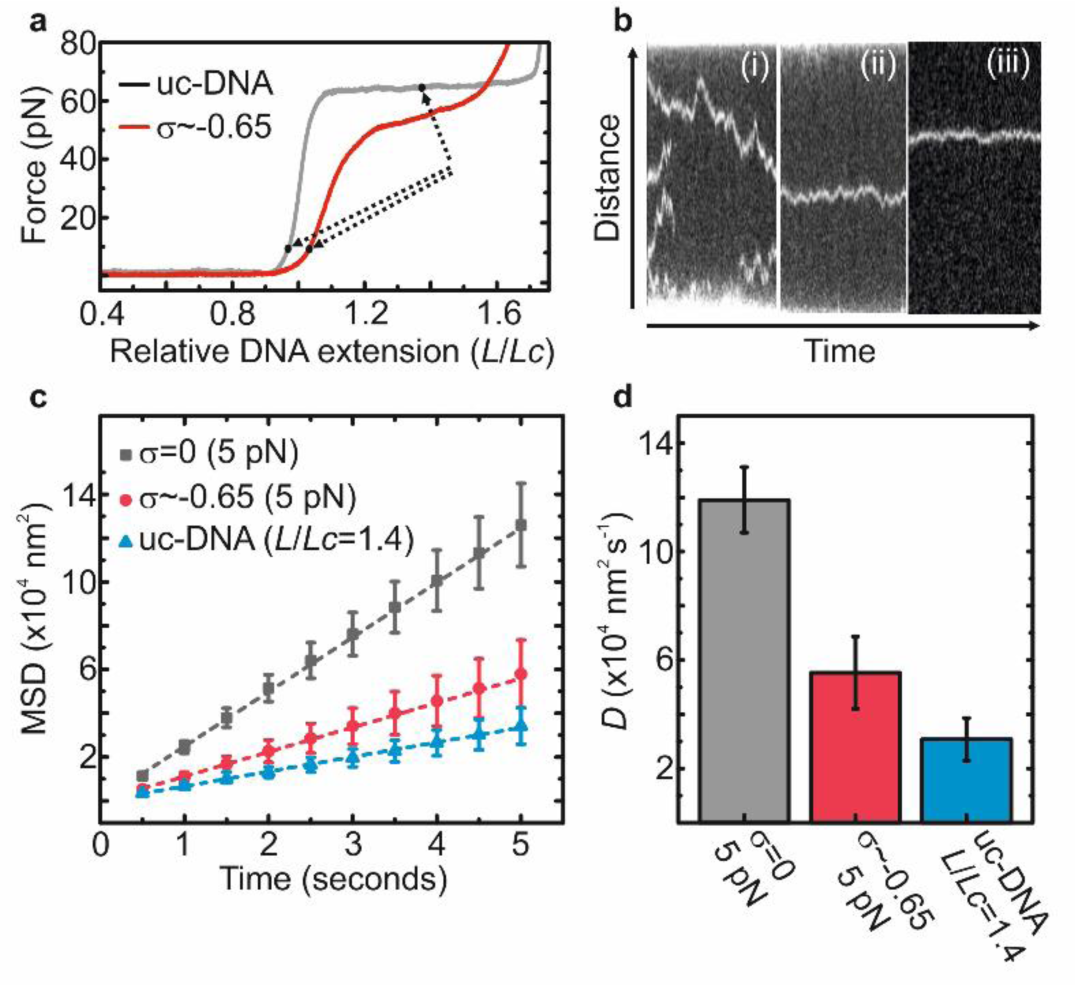
Application of ODS to study protein diffusion on negatively supercoiled DNA. **a**. Sample force-extension curves of end-closed torsionally unconstrained (uc) DNA (grey) and negatively supercoiled DNA (σ ~-0.65, red). Arrows highlight the conditions under which TFAM diffusion experiments were performed. **b**. Sample kymographs for alexa-555-TFAM monomers on (i) B-DNA at 5 pN and (ii) negatively supercoiled DNA (σ ~-0.65) at 5 pN and (iii) torsionally unconstrained DNA at *L*/*Lc* = 1.4. **c**. Mean squared displacement (MSD) of alexa-555-TFAM monomers as a function of time on B-DNA at 5 pN (grey), negatively supercoiled DNA (σ ~-0.65) at 5 pN (red) and torsionally unconstrained (uc-)DNA in an overstretched state (with relative contour length, *L*/*Lc* = 1.4; blue). Data points correspond to ~20 monomers (from ~10 independent DNA molecules) for each substrate. Dashed lines indicate linear fits to the data. **d**. Comparison of the diffusion coefficients (*D*) for alexa-555-TFAM monomers on B-DNA at 5 pN, negatively supercoiled DNA (σ ~-0.65) at 5 pN, and torsionally unconstrained (uc-)DNA at *L*/*Lc* = 1.4, respectively, extracted from the fits to the data in panel c. Note that no difference in the diffusion constant of TFAM was found when comparing torsionally constrained and torsionally unconstrained DNA at 5 pN (Fig. S9). All data were obtained in a buffer of 20 mM Tris-HCl, pH 7.6 with 25 mM NaCl. Errors are s.e.m.

While the mobility of the protein on non-supercoiled (B-form) DNA is rapid and minimally force dependent (Fig. S9), it is still lower than the maximum limit for rotation-coupled diffusion along the DNA backbone.^28^ This implies that the protein is slowed down by more stable transient interactions with the DNA. The current work suggests that these interactions are either stronger and / or less transient when the DNA is underwound. This is consistent with a recent study showing that TFAM has a higher binding affinity for negatively supercoiled plasmids.^29^ We thus propose that TFAM interacts more strongly with local underwound structures, resulting in a slower 1D sliding on DNA. This has potentially important implications for the ability of TFAM to locate and interact with promoter sites *in vivo* and suggests a mechanism by which supercoiling could regulate processes such as mitochondrial transcription.

Taken together, the applications presented in this study showcase the unique ability of ODS to interrogate long molecules of supercoiled DNA using a combination of 3D nano-manipulation and fluorescence microscopy. In this way, we demonstrate that the method is a robust and powerful approach for probing the structural properties of underwound DNA as well as the interplay between negative supercoiling and DNA-binding proteins. We propose that ODS can therefore be applied to address many open questions in genome biology, by enabling mechanistic studies of, for example, RNA polymerases, helicases and topoisomerases. To this end, we envision that the technique could be even further functionalized through integration with, for example, FRET,^24^ super-resolution imaging^23^ and fluorescence polarization microscopy.^30^

## Methods

### DNA construct design

End-closed DNA molecules (~48.5 kb) were prepared by ligating an end-cap to each terminal *cos*-site of λ-DNA.^18^ Each end-cap was composed of a 5T loop adjoined to a 12 bp double-stranded stem and a 12 nucleotide single-stranded overhang. The sequences of these end-caps are as follows:

End cap #1:

5’ AGG TCG CCG CCC GGA GTT GAA CG<u>T T</u>T<u>T</u>T<u>T</u> ACG TTC AAC TCC 3’

End cap #2:

5’ GGG CGG CGA CCT CAA GTT GGA CAA <u>T</u>T<u>T</u>T<u>T T</u>TG TCC AAC TTG 3’

Biotin moieties were covalently attached to four thymine residues (underlined above) within or near the 5T loop. Once each end-cap was ligated to λ-DNA, the DNA molecule could be tethered between two streptavidin-coated beads via biotin-streptavidin bonds. The number of biotins which bind to a given bead is variable: in the majority of cases (~90-95%), at least two biotins on each end of the DNA molecule bind to a bead. This renders the DNA molecule unable to change its total linking number and the molecule is thus torsionally constrained.

### Single-molecule assays

We employed a custom-built inverted microscope that combines dual-trap optical tweezers and wide-field fluorescence microscopy.^18, 20^ Experiments were performed in a multi-channel laminar flow-cell where end-closed DNA was tethered between two microspheres (each of diameter 4.5 μm) *in situ*. Such a flow-cell also allowed this dumbbell construct to be exchanged rapidly between channels containing different buffers or ligands. Forces applied to the DNA (via displacement of a tethered microsphere) were measured via back-focal plane interferometry of the condenser top lens using a position-sensitive detector.^23^ Data were obtained at room temperature in a buffer of 20 mM Tris-HCl, pH 7.6. The buffer also contained 25 mM NaCl, unless stated otherwise. eGFP-RPA and alexa-555-TFAM were prepared as detailed previously.^20, 28^

### Generation of negatively supercoiled DNA

The method employs the following protocol to generate negative supercoiling in DNA (see also Fig. 1a). (i) A torsionally constrained DNA substrate is prepared (as above) by tethering both strands of an end-capped DNA molecule to optically trapped microspheres. (ii) Tension is applied to the torsionally constrained DNA. The applied tension must be sufficient to induce overstretching. In torsionally constrained DNA, overstretching typically occurs at forces of ~115 pN.^18, 19^ Due to the torsional constraint, the *overall* linking number remains unchanged. However, overstretching is permitted in this case because some domains of the DNA molecule underwind, while other domains overwind.^18, 19^ For a fully overstretched torsionally constrained DNA molecule, ~4/5 of the molecule’s length is underwound (~37.5 bp/turn) and ~1/5 is overwound (~2.5 bp/turn).^19^ In this way, the DNA molecule can extend by up to ~70% without any change in overall linking number. (iii) Torsional constraint is transiently broken. In the absence of torsional constraint, overstretching of end-closed DNA is associated *only* with underwinding of the double-helix (~37.5 bp/turn). Therefore, relative to DNA that is torsionally unconstrained, overstretching of torsionally constrained DNA induces a torsional stress in the molecule. This torsional stress can be, at least partly, released by breaking a sufficient number of biotin-streptavidin tethers, such that only one tether exists on at least one end of the DNA molecule (allowing the DNA to swivel around the single tether). (iv) Once the torsional stress has been released, there is a probability that the broken tether reforms. If this occurs, the DNA molecule is once again torsionally constrained; however, now it is constrained with a lower linking number than that which it started with. (v) The tension is released; the DNA molecule is preserved in a negatively supercoiled state (with an overall linking number that is lower than that of B-DNA), even at low forces.

### Calibrating the extent of supercoiling

The fixed change in DNA linking number generated using ODS can be calibrated using reference force-extension curves. Such reference curves can be obtained either from literature or independently using alternative single-molecule approaches (such as magnetic or micro-pipette tweezers). In our case, we chose to compare our force-extension curves with those of Léger *at al*.^19^ To this end, we plotted the DNA extension at 70 pN as a function of σ. At this force, the reference data show a near linear relationship between DNA extension and σ (over the range of −0.1 < σ < −0.7; see Fig. 2b). By comparing the DNA extension from our measurements at 70 pN with those of the reference data, the value of σ generated in our method could be determined.

### TFAM diffusion analysis

End-closed DNA molecules were incubated in a low concentration of alexa-555-labelled TFAM (~5 nM) such that only a few protein monomers were bound. The DNA molecules were then transferred to a protein-free buffer channel where fluorescence movies were recorded with a frame rate of 0.5 s. Kymographs of these movies were then analyzed with a custom-written MATLAB-based program that tracked the position of each alexa-555-TFAM monomer on the DNA as a function of time. Only traces spanning longer than 10 s and which did not cross another were considered. The diffusion constant *(D*) was determined from MSD plots (MSD = 2*Dt* + offset) for all trajectories measured.

## Acknowledgements

We thank Mauro Modesti and Carolyn Suzuki for the kind gift of eGFP-RPA and TFAM, respectively. The authors are grateful to Andreas Biebricher for helpful discussions and Iddo Heller for providing kymograph position tracking software. The work was supported by funding through a Human Frontier Science Program grant (G.J.L.W), VICI grants from the Netherlands Organization for Scientific Research (NWO) (G.J.L.W and E.J.G.P), an NWO Chemical Sciences Top grant (G.J.L.W and E.J.G.P) and a long-term postdoctoral fellowship from the European Molecular Biology Organization (G.A.K).

