## Supplementary information for "Supercoiling DNA optically"

**Contents**

**Fig. S1.** Comparison of force-distance behaviours observed for end-capped DNA

**Fig. S2.** Proposed model for DNA-bead tethering geometries

**Fig. S3.** Predicted timescales for rotational motion of an optically trapped bead

**Fig. S4.** Temporal stability of the negatively supercoiled state

**Fig. S5.** Effect of ionic strength, laser power and stretching rate on underwound DNA

**Fig. S6.** eWLC fits to force-distance curves of negatively supercoiled DNA

**Fig. S7.** Force-extension curves of negatively supercoiled DNA in 0.8 nM and 5 nM RPA

**Fig. S8.** Kymograph of negatively supercoiled DNA in 0.8 nM eGFP-RPA

**Fig. S9.** Diffusion coefficient for alexa-555-TFAM on non-supercoiled DNA substrates.

**Supplementary Note 1.** Force-distance behaviours of end-capped DNA

**Supplementary Note 2.** Rotational velocity of optically trapped beads

**Supplementary Note 3.** Temporal stability of the supercoiled state

**Supplementary Note 4.** Structural analysis of underwound DNA

**Supplementary Note 5.** Influence of RPA on DNA structure

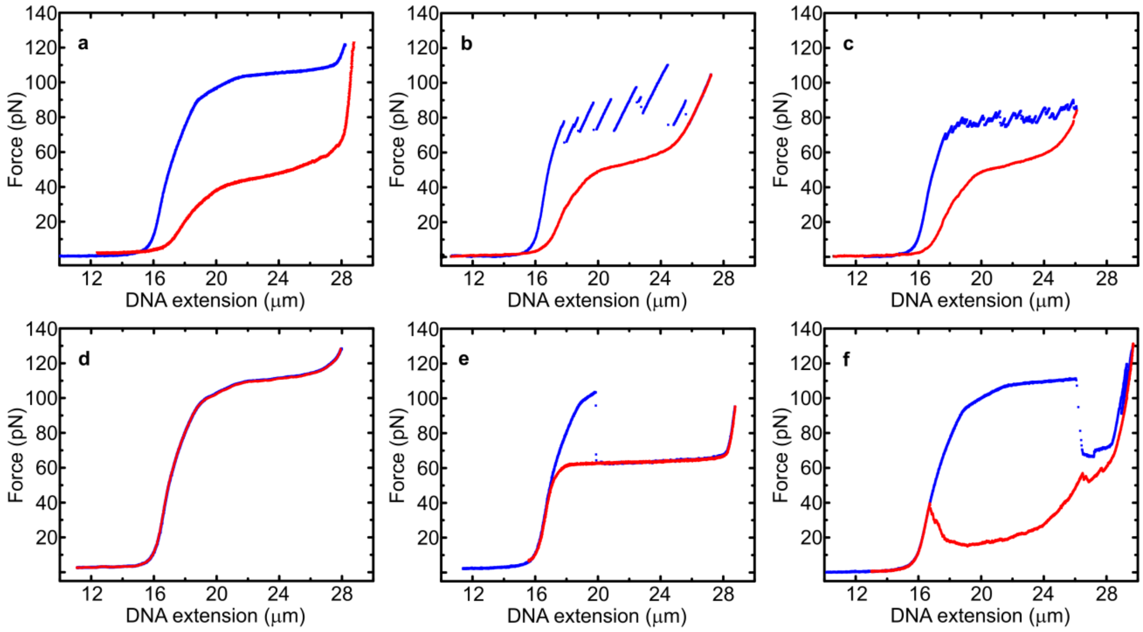


**Figure S1: Comparison of force-distance (FD) behaviours observed for end-capped DNA containing multiple biotin moieties.** In each case, the molecule begins in a torsionally constrained non-supercoiled state (σ = 0). A force-extension curve (blue) is recorded followed by a retraction curve (red). In this regard, six different behaviours are identified (Types 1-6). **a**. Type 1: supercoiled DNA (red) is observed only after initial extension to forces > 115 pN. The overstretching plateau associated with the initial FD curve (blue) is smooth. **b**. Type 2: the initial FD curve (blue) displays saw-tooth-like force ruptures at ~80-100 pN. After each jump in force, the DNA is increasingly supercoiled (red). **c**. Type 3: the initial FD curve (blue) displays a rugged force plateau at ~80 pN with many small force ruptures, which each give rise to increasing amounts of supercoiling (red). For both Type 2 and Type 3 behaviour, supercoiling is generated throughout the overstretching transition; the magnitude of σ formed in these cases correlates with the relative extension at which the force ruptures occur (Fig. 2e). **d**. Type 4: the DNA remains non-supercoiled (σ = 0), even when stretched to high forces (> 115 pN). Here the extension (blue) and retraction (red) curves are identical. **e**. Type 5: the initial torsionally constrained molecule (σ = 0) converts to unconstrained DNA via permanent cleavage of a biotin-streptavidin tether. In this case, the molecule is still end-closed, and thus exhibits a smooth overstretching plateau at ~65 pN.[1,2,3] **f**. Type 6: the initial torsionally constrained molecule ($\sigma = 0$) converts to unconstrained DNA via the (accidental) creation of a nick in the DNA backbone. The retraction curve (red) displays extensive hysteresis (at least under low salt conditions) due to strand-separation (peeling).[1,2] All data were obtained in a buffer containing 20 mM Tris-HCl, pH 7.6 with 25 mM NaCl.

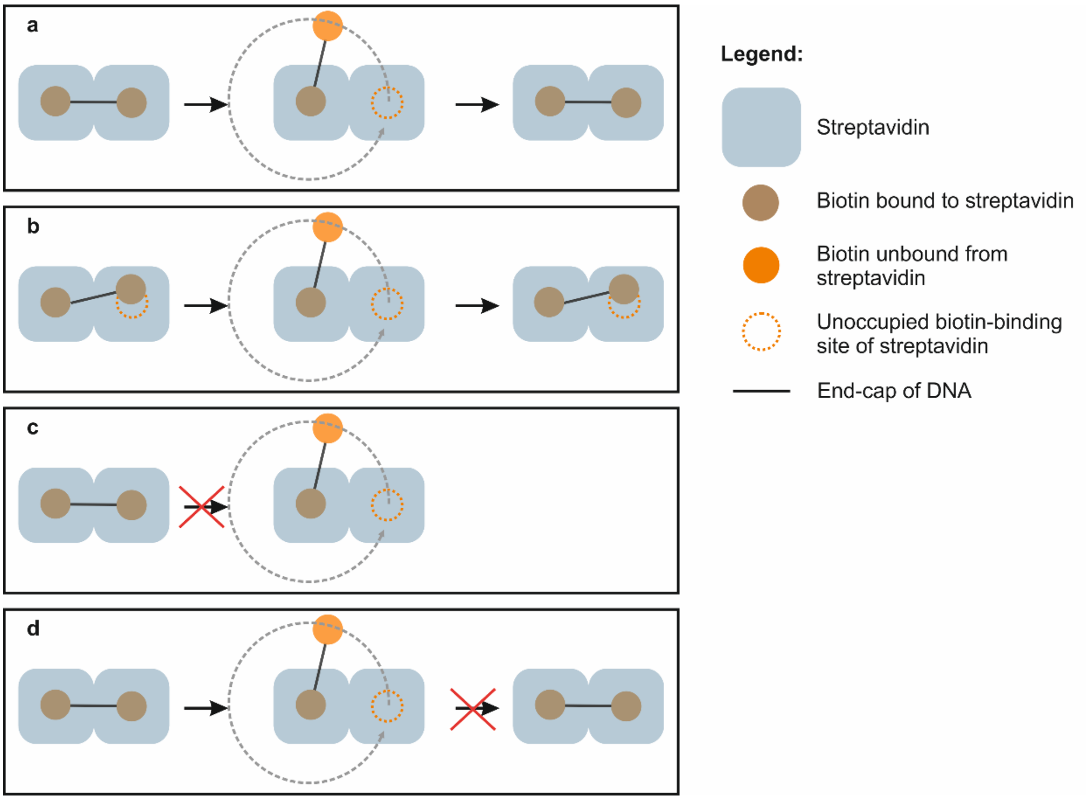


**Figure S2: Proposed model for how end-capped torsionally constrained DNA responds to high force.** Each end-cap is labelled with two biotin moieties that are initially tethered to a bead via interactions with streptavidin**. a**. One biotin dissociates from its streptavidin binding partner, allowing the DNA molecule to unwind by swiveling around the sole remaining biotin-streptavidin tether (represented by the dashed arrow). The disrupted bond subsequently reforms after DNA unwinding, resulting in negatively supercoiled DNA. This pathway typically occurs at forces > 115 pN and gives rise to the behaviour defined as Type 1 in Fig. S1a. **b**. One of the biotin moieties is initially weakly bound to a streptavidin partner (represented here by a partial overlap between the biotin and streptavidin). This bond is continuously broken and reformed as the molecule is progressively overstretched (at forces > 80 pN), resulting in the stick-slip force-extension pattern characteristic of Type 2 and Type 3 behaviour (as defined in Fig. S1b,c). In this case, each transient disruption of the tether results in the generation of supercoiling; the total amount of supercoiling here is proportional to the relative extension of the overstretched molecule (Fig. 2e). **c**. Both biotins on each end-cap of the DNA are stably bound to streptavidin, and do not dissociate, even under forces > 115 pN. The DNA is maintained in a non-supercoiled torsionally constrained state at all tensions (defined as Type 4 in Fig. S1d). **d**. One biotin dissociates from its streptavidin binding partner, allowing the DNA molecule to unwind by swiveling around the sole remaining biotin-streptavidin tether. Unlike in panels a and b, however, the broken tether never reforms; the end-closed DNA thus remains torsionally relaxed (defined as Type 5 behaviour in Fig. S1e).

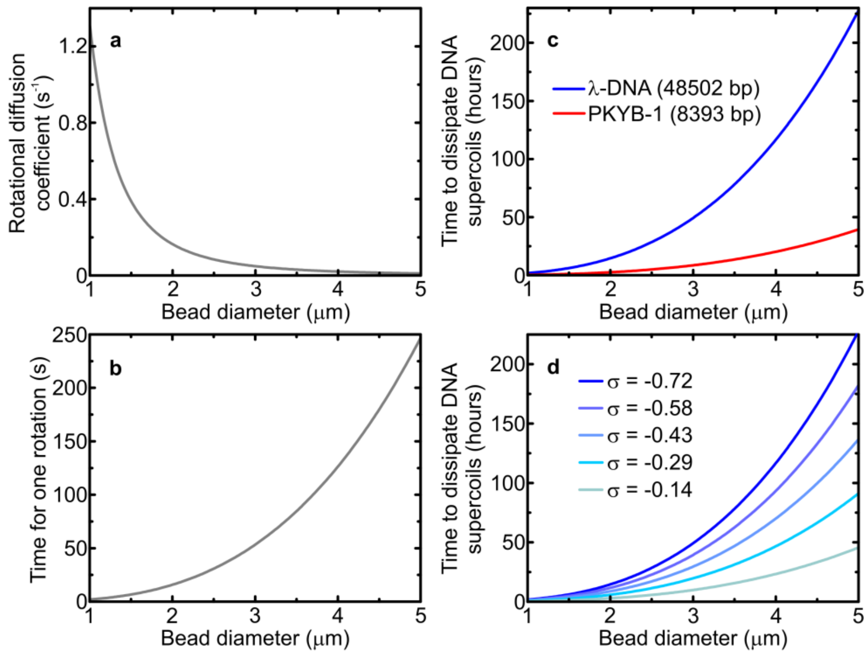


**Figure S3: Predicted timescales for rotational motion of an optically trapped bead. a**. Rotational diffusion coefficient of a spherical bead in water as a function of bead diameter, calculated using Eqn. 1 (SI Note 2). **b**. Predicted timescale for a single rotation of a spherical bead in water, when subject to an applied torque of -10 pN nm (determined using Eqn. 3, see SI Note 2). **c**. Predicted timescale for complete re-winding of long (λ-DNA) and short (PKYB-1) DNA constructs, as a function of bead diameter, assuming an initial state of $\sigma = -0.72$ (based on the data in panel b). **d**. Predicted timescale for complete re-winding of λ-DNA, as a function of bead diameter, for different initial extents of supercoiling ($-0.14 < \sigma < -0.72$) (based on the data in panel b).

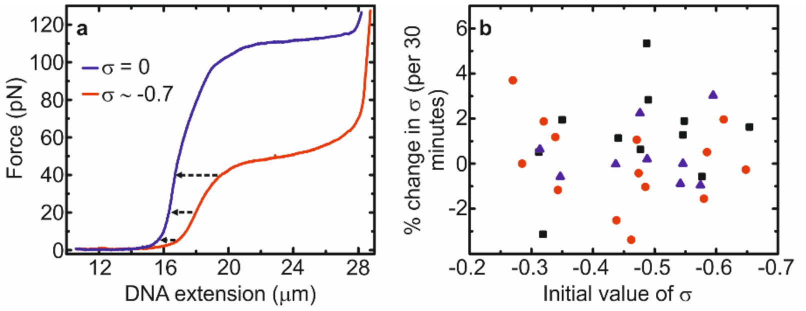


**Figure S4: Temporal stability of the negatively supercoiled state**. **a**. Experimental procedure: we measured the change in *Lk* (and thus σ) over time by monitoring the DNA extension at a constant force (5 pN, 20 pN and 40 pN, respectively). Since the supercoiled molecule is longer than non-supercoiled DNA at these forces, any change in σ over time is detected through a change in extension. As shown in Fig. 2f, a significant change in σ was only observed in a very small number of cases ($N = 3/30$). In the vast majority of cases, (90%; $N = 27/30$), negligible change in σ was detected over at least 30 minutes. **b**. Fractional change in σ associated with the data in the grey box in Fig. 2f, corresponding to the 90% of cases where negligible change in σ was detected (data collected at 5 pN (black), 20 pN (red) and 40 pN (blue)).

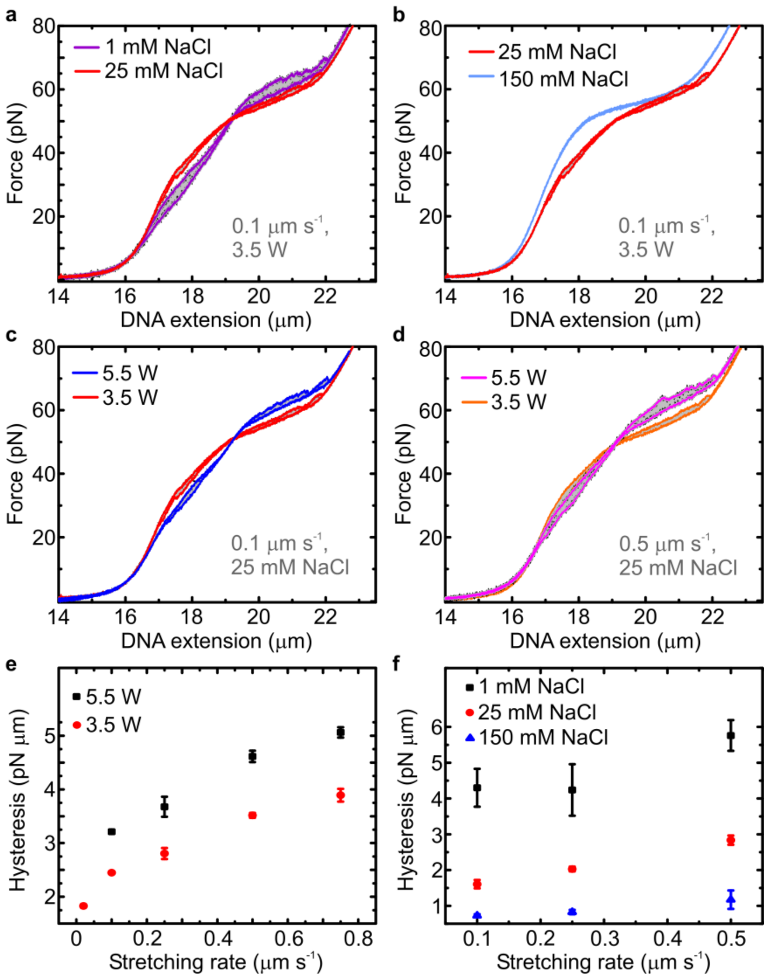


**Figure S5: Effect of ionic strength, laser power and stretching rate on the force-distance (FD) curves of negatively supercoiled DNA. a-b**. Effect of ionic strength on the measured FD curves at constant optical trapping laser power (3.5 W) and constant stretching rate (0.1 μm $s^{-1}$). **c**. Effect of the optical trapping laser power on the measured FD curves at constant ionic strength (25 mM NaCl) and constant stretching rate (0.1 μm $s^{-1}$). Note that the laser power will induce local heating of the buffer, and it thus serves as a proxy for temperature. **d**. Effect of the optical trapping laser power on the measured FD curves at constant ionic strength (25 mM NaCl) for a higher constant stretching rate than in panel c (0.5 μm $s^{-1}$). **e**. Measured hysteresis (determined from the area between extension and retraction FD curves) for two different laser powers as a function of stretching rate (at 25 mM NaCl). **f**. Measured hysteresis (determined from the area between extension and retraction FD curves) for different ionic strengths as a function of stretching rate (at 3.5 W laser power). All data were obtained for $\sigma$ ~-0.45, in a buffer containing 20 mM Tris-HCl, pH 7.6, with the relevant ionic strengths specified above. Errors are s.e.m. (N~5).

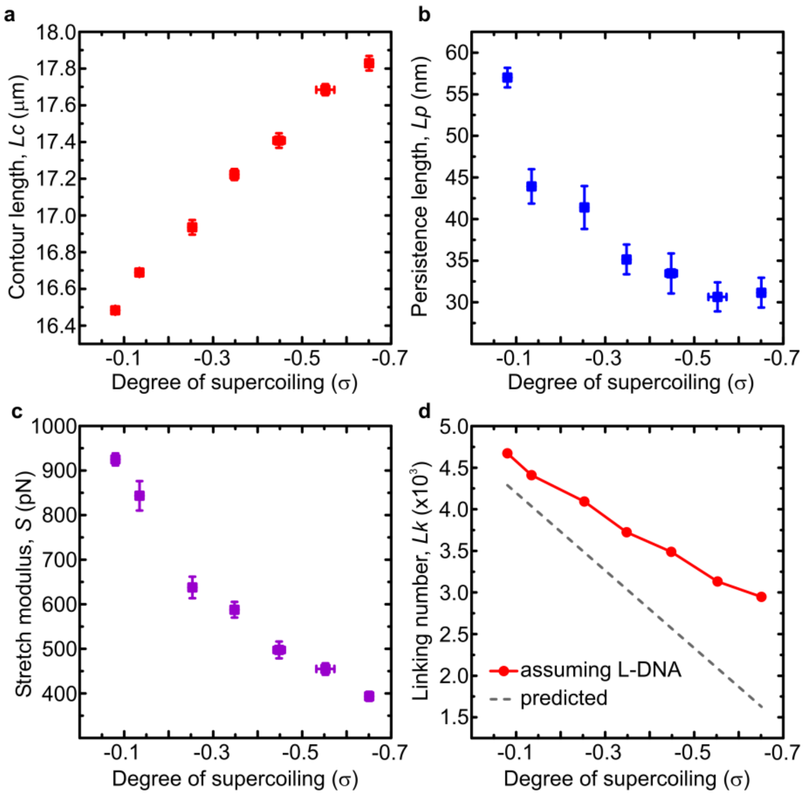


**Figure S6: Mechanical properties of underwound DNA as a function of negative supercoiling. a-c**. Results of fits of the extensible Worm-Like Chain (eWLC) model (up to 30 pN) to experimentally derived force-distance curves of end-closed DNA (*e.g*. Fig. 2a) as a function of supercoiling (σ). Note that the DNA is expected to consist of regions of both B-DNA and underwound structures (such as L-DNA). Thus, the contour length (*Lc*), persistence length (*Lp*) and stretch modulus (*S*) extracted from eWLC fits are average values for the entire DNA molecule, rather than for one specific structure. **d**. The red circles show the change in DNA linking number (*Lk*) as a function of σ, assuming that the difference in *Lc* between supercoiled and non-supercoiled DNA (identified from panel a) is due solely to the formation of L-DNA (SI Note 4). For comparison, the expected change in *Lk* as a function of σ is highlighted by the grey dashed line. As explained in SI Note 4, the observed difference here likely reflects the additional presence of more compact DNA structures (such as Z-DNA) that co-exist with B-DNA, L-DNA and bubble-melted DNA. All data (*N*~10 for each value of σ) were obtained in a buffer containing 20 mM Tris-HCl, pH 7.6 with 25 mM NaCl. Errors are s.e.m.

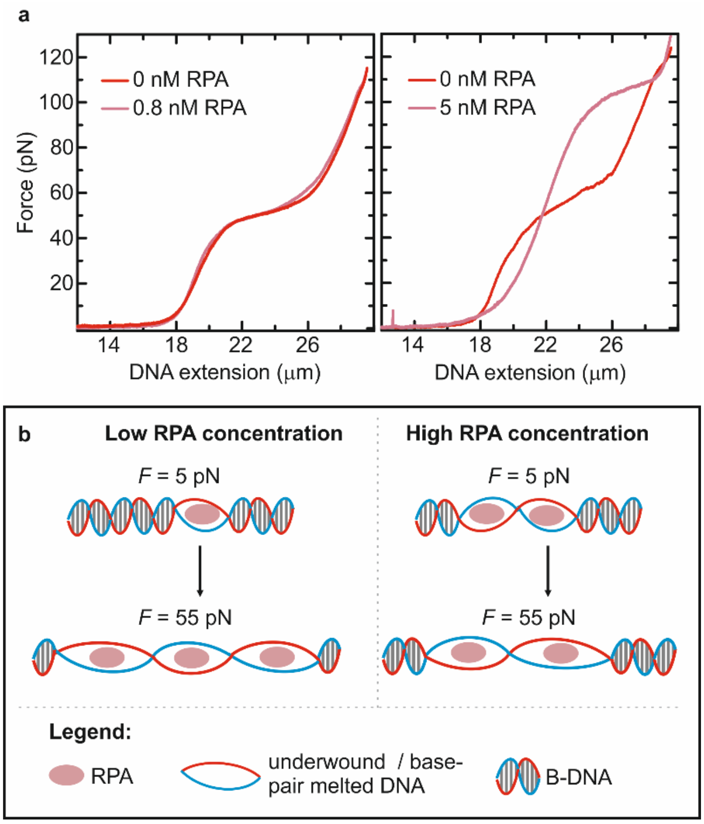


**Figure S7: Influence of RPA on the structure of negatively supercoiled DNA. a**. Comparison of force-extension curves of negatively supercoiled DNA (σ ~ -0.65) in the presence of low (0.8 nM) and high (5 nM) concentrations of eGFP-RPA. **b**. Proposed model for how RPA interacts with negatively supercoiled DNA. In low concentrations, RPA binds to base-pair melted structures, but does not significantly perturb the DNA. Thus, at low forces (*F* = 5 pN), small localized regions of bubble-melted DNA are bound by RPA, while at ~55 pN extensive RPA binding occurs (due to the formation of overstretched bubble-melted structures). In contrast, at high concentrations, RPA can create additional regions of bubble-melted DNA at 5 pN. These bubble-melted domains are more highly underwound than those that are favoured at 55 pN and are topologically 'locked' by the bound RPA. As a result, the remaining B-DNA domains present at 5 pN cannot unwind at 55 pN as they would normally do. Rather, these B-DNA regions will overstretch at much higher forces (~115 pN), as illustrated in the right-hand side of panel a. See SI Note 5 for more details. Data were obtained in a buffer containing 20 mM Tris-HCl, pH 7.6 with 25 mM NaCl.

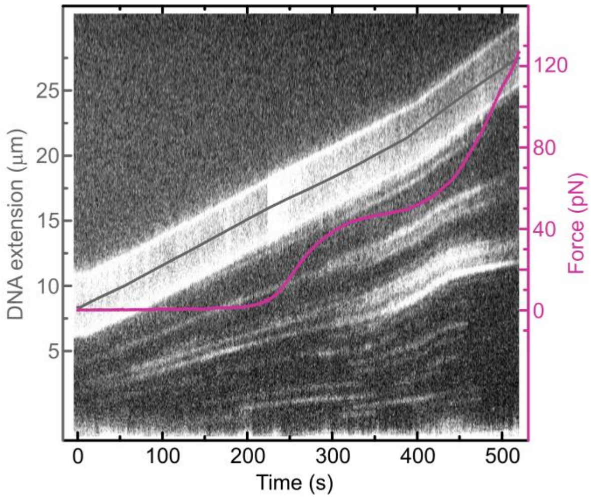


**Figure S8: Illustrative kymograph for stretching negatively supercoiled DNA (σ ~-0.65) in the presence of eGFP-RPA (0.8 nM).** Here, the DNA molecule is stretched from low force (< 1 pN) to high force (~120 pN). The measured DNA extension and force, as a function of time, are overlaid on the fluorescence image in grey and pink, respectively. A clear increase in fluorescence intensity is observed after the onset of the force plateau (at ~50-60 pN). This is consistent with our observations (Fig. 3) that more RPA binds to underwound DNA at ~55 pN compared with 5 pN. Note that the bright diagonal in the kymograph is the moving optically trapped bead. Data were obtained in a buffer containing 20 mM Tris-HCl, pH 7.6 with 25 mM NaCl.

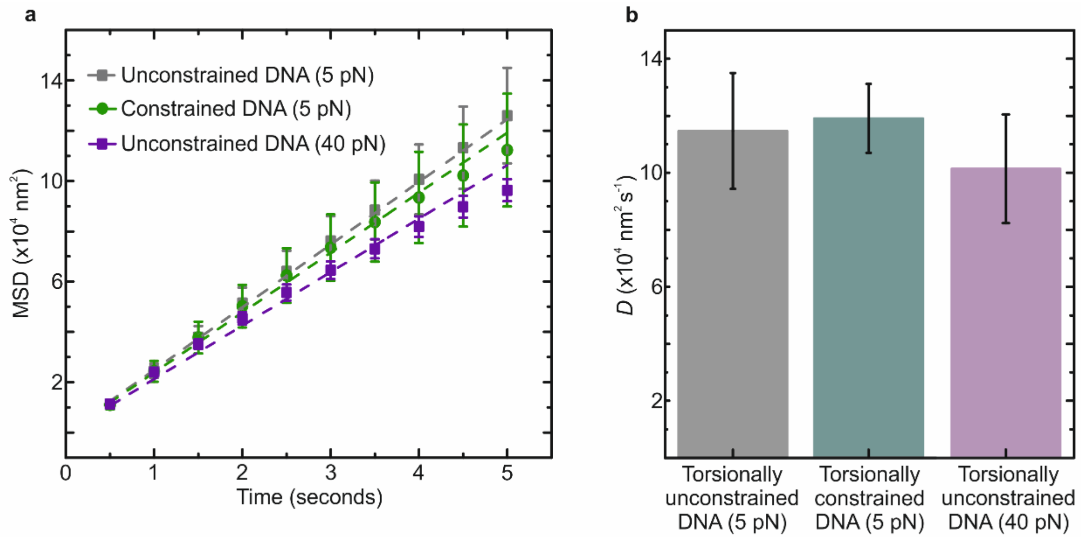


**Figure S9: Comparison of the diffusion coefficient for alexa-555-TFAM on different non-supercoiled DNA substrates. a**. Mean squared displacement (MSD) of alexa-555-TFAM monomers as a function of time on torsionally unconstrained DNA at 5 pN (grey), torsionally constrained DNA ($\sigma = 0$) at 5 pN (green) and torsionally unconstrained DNA at 40 pN (purple). Data points correspond to ~20 monomers (from ~10 independent DNA molecules) for each substrate. Dashed lines indicate linear fits to the data. **b**. Diffusion coefficients for monomers of alexa-555-TFAM on torsionally unconstrained DNA at 5 pN (grey), torsionally constrained DNA ($\sigma = 0$) at 5 pN (green) and torsionally unconstrained DNA at 40 pN (purple), extracted from fits to the data in panel a. All data were obtained in a buffer of 20 mM Tris-HCl, pH 7.6 with 25 mM NaCl. Errors are s.e.m.

**Supplementary Note 1: Force-distance behaviours of end-capped DNA**

The end-capped DNA construct used throughout this work can give rise to a number of different force-distance behaviours. Under the conditions employed in our experiments, we identified six principle types of behaviour, as detailed below (and illustrated in Fig. S1).

**Type 1:** Torsionally constrained DNA ($\sigma = 0$) is converted to a supercoiled state following extension to high forces (> 115 pN). This is characterized by a smooth initial overstretching transition of the torsionally constrained molecule (*e.g.* Fig. S1a), and constitutes ~25% of cases.

**Type 2:** Torsionally constrained molecules ($\sigma = 0$) are converted to a supercoiled state via abrupt force ruptures between 80 pN and 120 pN (*e.g.* Fig. S1b). This behaviour occurs in ~25% of cases.

**Type 3:** Torsionally constrained molecules ($\sigma = 0$) are converted to a supercoiled state via a rough force-extension plateau at ~80 pN. This occurs in ~5-10% of cases, and is exemplified in Fig. S1c.

**Type 4:** DNA molecules are stably preserved in a non-supercoiled torsionally constrained state even under high tension (> 200 pN) and cannot easily be converted to a supercoiled state. This behaviour (illustrated in Fig. S1d) occurs in ~5-10% of cases.

**Type 5:** DNA molecules switch from torsionally constrained ($\sigma = 0$) to unconstrained during the overstretching transition (Fig. S1e). This occurs due to permanent cleavage of a biotin-streptavidin tether. Since these molecules have no open ends or nicks (*i*.e. they are topologically closed), a smooth overstretching plateau at ~65 pN is observed. This behaviour occurs in ~5-10% of cases.

**Type 6:** DNA molecules contain at least one nick in their phosphate backbone, as a result of thermal, mechanical or chemical degradation, resulting in a lack of torsional constraint. Such nicks can occur in solution, prior to single-molecule manipulation or during manipulation (*e.g*. Fig. S1f). In our sample, ~25-30% of molecules were nicked.

**Supplementary Note 2: Rotational velocity of optically trapped beads due to applied torque**

The ability of ODS to produce and maintain a reduced DNA linking number relies, in part, on the assumption that the supercoiling generated is not easily dissipated through free rotation of the optically trapped beads. To validate this assumption, we first note that the rotational diffusion coefficient (*D*) is given by the Einstein relation:

$$\boldsymbol{D} = \frac{\boldsymbol{KT}}{\boldsymbol{f_r}} \qquad (1)$$

where KT is the thermal energy and $f_r$ is the frictional drag coefficient. The latter is defined as follows:

$$\boldsymbol{f_r} = \mathbf{8\pi\eta}\boldsymbol{R^3} \quad (2)$$

Here, *R* is the radius of the bead, and $\eta$ is the dynamic viscosity of the solution. Owing to its dependence on $R^3$, the rotational diffusion is reduced significantly when using larger beads (Fig. S3a). When tethered to a supercoiled DNA molecule, the beads will experience an applied torque (Γ). The rotational velocity (Ω) of the beads in response to an applied torque can be calculated using the following relation:

$$\Omega = \frac{\Gamma}{f_r} \qquad (3)$$

It has previously been reported that for supercoiled DNA in the range of $-0.1 < \sigma < -1.5$, the torque is roughly constant at approximately -10 pN nm.[4,5] Fig. S3b plots the predicted timescale for a single bead rotation (using Eqn. 3) as a result of such torque as a function of bead diameter. Note that the time is significantly longer (~180 s) when using large beads (*i.e.* 4.5 μm diameter) compared with smaller beads (*i.e.* 1 μm diameter). Moreover, since each rotation of the bead results in a unit change of the DNA linking number ($\Delta Lk$ = +1), the timescale for converting the entire molecule from a supercoiled state to a non-supercoiled state depends on two additional factors. The first is the total number of base-pairs in the molecule. As Fig S3c demonstrates, assuming a torque of -10 pN nm and 4.5 μm diameter beads, it should take ~170 hours to fully convert a λ-DNA molecule from a highly supercoiled state (σ = -0.72) to a non-supercoiled state through rotational motion of the beads. This contrasts with the case of PKYB-1 (~8 kb), which is expected to dissipate a similar degree of supercoiling in ~30 hours. A second factor is the extent of initial supercoiling: the fractional change in *Lk* over time is inversely proportional to the initial value of σ, as highlighted in Fig. S3d. Nonetheless, assuming long DNA and large beads are used, the total change in *Lk* should be minimal on the timescale of most single-molecule experiments. For example, when using 4.5 μm diameter beads, and λ-DNA, the fractional change in linking number over a period of 30 minutes is expected to range from ~0.3% (for an initial σ of -0.7) to ~1.5% (for an initial σ of -0.1). Finally, we note that any small asymmetries in the shape of the beads will tend to align the beads in a particular orientation in the optical traps; this would further be expected to hinder the rotational motion of the beads.

**Supplementary Note 3: Temporal stability of the supercoiled state**

The temporal stability of the supercoiled state generated using ODS is affected by two main factors. The first is the speed at which torsional stress can be relieved through rotational motion of the tethered beads. The calculations detailed in SI Note 2 indicate that for long DNA molecules and large beads, this speed is small (< 1.5% change in *Lk* over 30 minutes). The second factor affecting the lifetime of the supercoiled state is the intrinsic stability of the biotin-streptavidin bonds: any transient loss of torsional constraint due to cleavage of a tether could lead to rewinding of the double-helix and loss of supercoiling. Indeed, it is the transient cleavage of a biotin-streptavidin tether that underpins the principle of ODS. However, such cleavage is promoted by tension; it has been predicted previously that the lifetime of a biotin-streptavidin bond is several orders of magnitude lower at forces > 100 pN than at forces < 10 pN.[6] Taken together, this suggests that, once generated, the supercoiled state should be stable over significant periods of time (at least many tens of minutes).

To test the above assertions, we measured the change in *Lk* (and thus $\sigma$) over time by monitoring the DNA extension at a fixed force (5 pN, 20 pN and 40 pN, respectively). At these tensions, negatively supercoiled DNA (for $\sigma$ in the range of -0.1 to -0.7) is longer than for B-form DNA, and thus any change in $\sigma$ should be detected through a change in extension (Fig. S4a). For the vast majority of molecules considered (*N* = 27/30), negligible change in extension (and thus $\sigma$) was detected over a 30-minute period. However, in ~10% of cases (*N* = 3/30) a significant change in $\sigma$ (between 20% and 100%) was observed within this time span. We note that all unstable molecules were characterized as having Type 3 behaviour (as defined in SI Note 1 and Fig. S1c). In contrast, Type 1 and Type 2 molecules typically lead to stable supercoiled states at forces < 40 pN over long (at least 30 minute) durations (as do some Type 3 molecules).

**Supplementary Note 4: Structural analysis of underwound DNA**

It is well-established that, upon tuning σ from -0.1 to -1.5, torsionally constrained DNA held at ~5 pN undergoes a cooperative transition from B-DNA to an underwound state. During this transition the torque remains constant at ~-10 pN nm.[4,5] The structure of the resulting underwound state is attributed largely to a left-handed conformation known as L-DNA. L-DNA has been estimated to exhibit a helicity of ~-12 bp/turn, a persistence length of 3 nm and a contour length of 0.48 nm/bp.[4] Nonetheless, there is mounting evidence that negatively supercoiled DNA can adopt additional conformations during the unwinding transition.[4,5,7,8,9] These include bubble-melted DNA (M-DNA) and Z-DNA The former refers to a base-pair melted underwound structure, while the latter is a left-handed, base-paired conformation with a similar helicity to L-DNA.[7] However, the exact structure of L-DNA and M-DNA in supercoiled DNA remains unclear, as does the role of the local environment in tuning their relative stabilities. Our results shed new light on these structures through a combination of force-spectroscopy and fluorescence microscopy, as detailed below.

**Worm-like chain analysis.** Force-extension curves of DNA were fit to the eWLC model (up to 30 pN) using Eqn. 4:[10]

$$\boldsymbol{d = L_{\mathrm{c}}\left(1 - \frac{1}{2}\left(\frac{k_{\mathrm{B}}T}{FL_{\mathrm{p}}}\right)^{1/2} + \frac{F}{S}\right)} \qquad (4)$$

Here, $d$ is the DNA extension at a given force $F$ and $k_B$ is the Boltzmann constant. $Lc$, $Lp$ and $S$ are the contour length, persistence length and stretch modulus of the molecule, respectively. By fitting force-extension curves of DNA to Eqn. 4 as a function of σ, we plot how supercoiling influences the global contour length, persistence length and stretch modulus of the molecule (Fig. 3c and Fig. S6a-c). We stress that these parameters are average values, and reflect contributions from each of the distinct structures that may co-exist within the molecule. Nonetheless, in this way, we deduce that, for increasingly negative supercoiling, the average value of $Lc$ increases, while the average values of $Lp$ and $S$ decrease. These indicate that underwound DNA is more extended and more flexible than B-form DNA. Using the relation $\sigma = (Lk\text{-}Lk_0)/Lk_0$, the change in $Lk$ as a function of $\sigma$ for λ-DNA is displayed by the grey line in Fig. S6d. We can compare this theoretical change in $Lk$ with that deduced from our eWLC fits. For example, if we assume that the change in average contour length for each value of $\sigma$ (as documented in Fig. S6a) is due to the formation of only L-DNA (alongside regions of B-DNA) we can calculate the corresponding change in $Lk$ that would be expected as a function of $\sigma$. As shown by the red data in Fig. S6d, this calculation does not fully agree with the expected relation, especially for increasingly negative values of $\sigma$. This could be accounted for if local regions of Z-DNA (a highly compact underwound structure) can exist alongside L-DNA, M-DNA and B-DNA. This is consistent with recent magnetic tweezers studies that reported the presence of Z-DNA in negatively supercoiled molecules.[5]

**Force-extension analysis.** If M-DNA can co-exist in negatively supercoiled DNA at low tensions, we anticipate that it will be more likely at lower salt concentrations (which favour base-pair melting).[1,2] This is consistent with the force-extension data presented in Fig. 3a,b and Fig. S5. For instance, in Fig. 3a, we observe a slight decrease in the apparent persistence length of negatively supercoiled DNA at low NaCl concentrations (< 25 mM). Second, the overstretching transition appears significantly less cooperative in lower salt concentrations, compared with higher ionic strengths, suggesting that the DNA molecule exists in a heterogeneous state. Finally, a small but clear hysteresis is observed when comparing extension and retraction force-distance curves at very low salt concentrations

(Fig. 3b) and also at high laser powers / temperatures (Fig. S5c,d). A similar hysteresis has been observed following overstretching of non-supercoiled DNA and attributed to base-pair melting.[1,2] For all of these reasons, we argue that some base-pair melting occurs in negatively supercoiled DNA at low forces (*e.g.* 5 pN), especially at low ionic strength and high temperature.

**eGFP-RPA binding.** Fluorescently-labelled RPA has been used previously to detect base-pair melted DNA generated via both tension and torque.[1,4,11] The protein has a footprint of around 30 nucleotides and binds to single-stranded DNA in a tension-independent manner.[1] Our analysis shows that, at low RPA concentrations and low salt concentrations, there is approximately a 3-4-fold higher RPA binding to supercoiled DNA ($\sigma$ ~-0.65) at 55 pN (where the molecule is overstretched) than at 5 pN (Fig. 3e). This confirms that the overstretched supercoiled molecule largely consists of extended bubble-melted DNA (at low ionic strength). However, the presence of some RPA binding at 5 pN substantiates our earlier conclusions (based on force-extension analysis) that some form of base-pair melted DNA can also occur at low forces.

**Supplementary Note 5: Influence of RPA on DNA structure**

It has been demonstrated previously that RPA can induce unwinding of the DNA double-helix. For instance, for negatively supercoiled DNA at very low tensions (< 1 pN), RPA can induce plectonemes to convert to bubble-melted conformations.[12] Similarly, for overstretched unconstrained DNA, RPA can increase the ratio of strand separation to base-paired S-DNA.[1] RPA has also been shown to catalyze the opening of DNA hairpins.[13] Nonetheless, previous work has also confirmed that, for low RPA concentrations, the perturbation to DNA structure is limited and, under these conditions, the protein effectively serves as a reporter for bubble-melted and single-stranded DNA.[1]

In the current work, we found that the perturbation to the mechanical properties of negatively supercoiled DNA was small for RPA concentrations of 0.8 nM, but significant for RPA concentrations of 5 nM (Fig. S7).

We note that at low RPA concentrations (0.8 nM), the protein only minimally perturbs the structure of negatively supercoiled DNA. To highlight this, Fig. S7a demonstrates that the force-extension curves of negatively supercoiled DNA in the presence of 0.8 nM RPA are very similar to those recorded in the absence of RPA. However, at higher RPA concentrations (5 nM), two major changes in the force-extension behaviour are observed. First, the negatively supercoiled molecule exhibits a longer contour length, indicating a more extended state at low force. Second, the force plateau (normally present at ~55 pN) now occurs at ~115 pN - similar to that of non-supercoiled torsionally constrained DNA.[11] The most likely explanation for these observations is that RPA ‘locks’ the supercoiled molecule in a highly underwound state at low tensions. In the absence of RPA, local regions of B-DNA (which co-exist alongside underwound structures in supercoiled DNA at low force) convert to an extended underwound state at ~55 pN. This can be achieved through a redistribution of twist throughout the molecule; namely, the most highly underwound regions present at low force become slightly less underwound at high tension, thereby allowing the B-DNA domains to adopt an underwound structure also. Our data indicate that RPA prevents this redistribution of twist, and thus the B-DNA domains will overstretch in a similar fashion to that of non-supercoiled torsionally constrained DNA (illustrated schematically in Fig. S7b). In this regard, our results support previous studies which have shown RPA can stabilize underwound DNA structures.[14,15]
